# A segmentation scheme for complex neuronal arbors and application to vibration sensitive neurons in the honeybee brain

**DOI:** 10.1101/323758

**Authors:** Hidetoshi Ikeno, Ajayrama Kumaraswamy, Kazuki Kai, Thomas Wachtler, Hiroyuki Ai

**Author notes:** **Correspondence:** Hidetoshi Ikeno. **Funding** This research was supported by Ministry of Education, Science, Technology, Sports and Culture of Japan, Grant-in-Aids for Scientific Research (C), grant No. 25330342, for Challenging Exploratory Research (15K14569) and Strategic International Cooperative Program, Japan Science and Technology Agency (JST) and the Federal Ministry of Education and Research of Germany (BMBF, grant 01GQ1116).

## Abstract

The morphology of a neuron is strongly related to its physiological properties, and thus to information processing functions. Optical microscope images are widely used for extracting the structure of neurons. Although several approaches have been proposed to trace and extract complex neuronal structures from microscopy images, available methods remain prone to errors. In this study, we present a practical scheme for processing confocal microscope images and reconstructing neuronal structures. We evaluated this scheme using image data samples and associated gold standard reconstructions from the BigNeuron Project. In these samples, dendritic arbors belonging to multiple projection branches of the same neuron overlapped in space, making it difficult to automatically and accurately trace their structural connectivity. Our proposed scheme, which combines several software tools for image masking and filtering with an existing tool for dendritic segmentation and tracing, outperformed state-of-the-art automatic methods in reconstructing such neuron structures. For evaluating our scheme, we applied it to a honeybee local interneuron, DL-Int-1, which has complex arbors and is considered to be a critical neuron for encoding the information indicated in the waggle dance of the honeybee.

## 1 Introduction

Neuronal morphology is strongly related to neuronal function; neuron arborization reflects the input and output regions, and morphological characteristics, such as branching pattern, length and thickness, are related to signal transmission properties. Changes of neuronal morphology have been observed in various nervous systems, especially as a consequence of development or experience (Jan and Jan, 2003, Grueber et al., 2005, Yasunaga et al., 2010, Luebke et al., 2015). Neuron segmentation and modeling is an effective way for quantitatively evaluating morphological properties of neurons and can be shared among researchers as a common resource (Halavi et al., 2008). Neuron morphology is most commonly extracted from image stacks that are obtained by scanning dye-filled brain tissues under confocal laser scanning microscopes (CLSM). To extract and trace neuronal structure from confocal images, we have used the software SIGEN (Yamasaki et al., 2006, Minemoto et al., 2009) and applied it for segmentation of neurons in the insect brain. This program uses a simple algorithm called single seed distance transform, which works effectively for segmentation of various types of neurons. We have previously used SIGEN to reconstruct several neurons from the moth brain, which were collected into a database and used for neuron morphological analyses and network simulations (Ikeno et al., 2012).

Automatic segmentation of neuronal structure from CLSM images is one of the most challenging themes in biological image processing, and several projects encompassing multiple research labs have been undertaken to foster better algorithms (Acciai et al., 2016). The DIADEM challenge (Brown et al., 2011) was a competition held to raise awareness of this problem and spur development of automatic reconstruction algorithms. In this project, a metric program for comparing two digital morphological reconstructions was developed (Gillette et al., 2011) and continues to be a widely used measure. Recently, the BigNeuron project has been conducted to develop automatic segmentation methods from confocal images (Peng et al., 2015). Although automatic neuronal segmentation and reconstruction has significantly advanced by these efforts, it is still difficult to correctly reconstruct complex structured neurons (Magliaro et al., 2017). Hence, it can be useful to combine automatic segmentation tools with manual operations for effective analysis of neuronal structure.

An interesting application for neural structure analysis is investigation of the neural mechanisms for encoding and decoding the spatial information to the profitable flower, indicated by honeybee waggle dance (von Frisch, 1967). Honeybee use a unique temporal pattern of vibration caused by wing beats for informing spatial information to their hive mates. Airborne vibrations are received by sensory neurons located in the Johnston’s organ of the antenna. These neurons mainly project their neurites into the dorsal lobes (DLs) in the honeybee brain. Among several types of interneurons with sensitivity to vibratory stimuli (Ai et al., 2009, Ai et al., 2017), DL-Int-1 is a major interneuron arborizing in the DL (Ai et al, 2009). This neuron shows inhibitory responses to vibratory stimuli applied to the antenna and has a unique structure. The soma is located close to the dorsal central body in the posterior protocerebral lobe, and a primary neurite extends into the DL. In the DL, two thick branches, called dorsal branch (DB) and ventral branch (VB), emerge from the primary neurite (Fig 5c; DB: branch towards n_2_, VB: branch towards n_3_). DB mainly arborizes on the dorsal side of the DL, whereas VB arborizes on the ventral side of the DL and extends into the subesophageal ganglion. These two branches overlap densely in the DL, which makes it difficult to separate them by any automatic segmentation process. To make a segmentation of the morphology of this interneuron, we developed and applied a practical scheme, which combines manual extraction of neural branches with application of automatic segmentation software to reconstruct neuron morphology. Our segmentation scheme can be applied to segment neurons with complex structures, especially when multiple projecting branches have dendritic arborizations in the same region. This scheme uses our original segmentation software, SIGEN (Minemoto, 2014) to extract and trace neurites automatically. We evaluate its performance before applying our new scheme to DL-Int-1.

**Figure 5.**
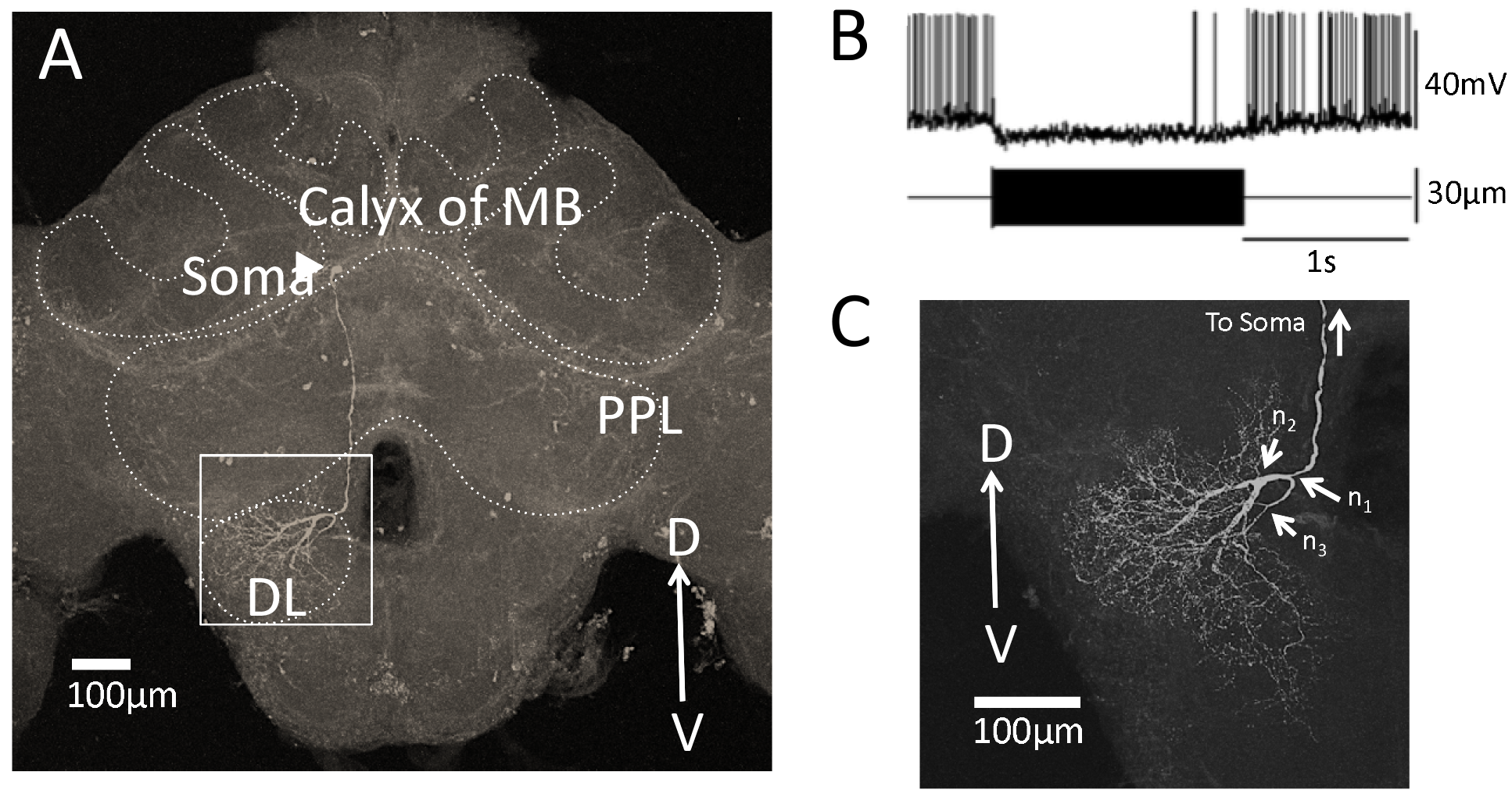
Segmentation of a vibration sensitive interneuron in the honeybee brain, DL-Int-1 **A.** The whole morphology of DL-Int-1 in the honeybee brain. The soma (arrow head) is located below the calyx of mushroom body. A long primary neurite extends ventrally and arborizes in the dorsal lobe. **B.** Typical unique electrophysiological response pattern of DL-Int-1. Tonic inhibitory response was observed by a continuous 265 Hz vibration stimulus to the antenna. The upper trace shows the action potentials of the DL-Int-1 (bottom trace: vibration stimulus). **C.** Arborization of neurites of DL-Int-1 in the dorsal lobe. n_1_ is the point at which the primary neurite bifurcated into the dorsal branch and ventral branch, which are dificult to spatially separate. The dorsal branch started from the branching point n_2_. The ventral branch started from the branching point n_3_.

## 2 Material and Methods

### 2.1 Neuronal segmentation and evaluation

Our scheme for segmentation of confocal image data starts with deconvolution by Fiji (RRID:SCR_002285) (Fig. 1A]). We used the “Iterative Deconvolve 3D” plugin along with the point spread function (PSF) that was generated by the “Diffraction PSF 3D” plugin with default parameter values (Dougherty, 2005). This is an effective process for decreasing blurring and distortion in confocal images. Next, we conducted neuron segmentation by SIGEN (RRID: SCR_016284), which was applied to the deconvolved images. Segmentation results were obtained in SWC format, which is widely used as a standard data format to describe neural structure (Ascoli et al., 2007).

**Figure 1.**
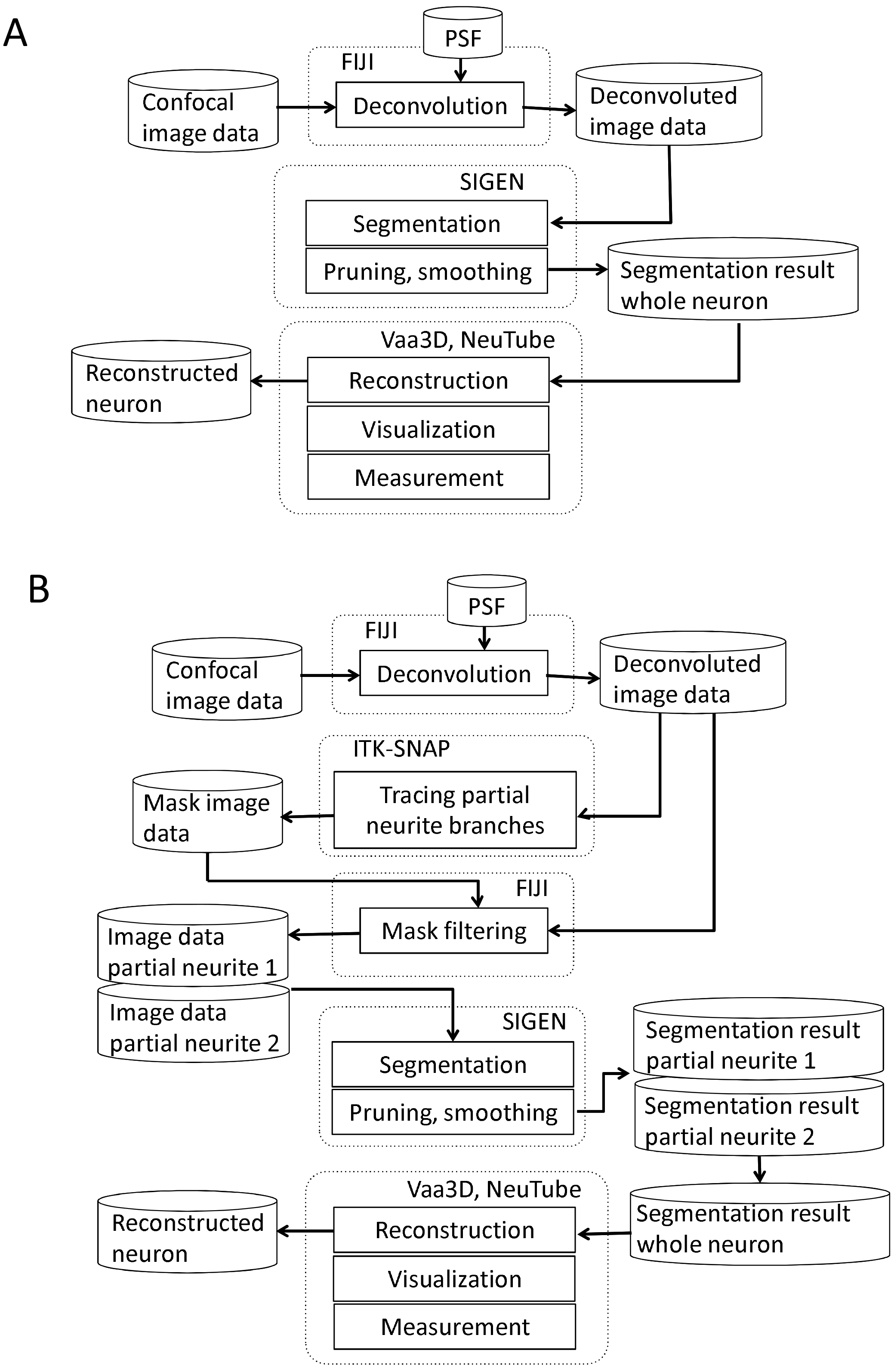
**A**. Our basic neuron segmentation scheme from a CLSM image. The CLSM image was deconvolved by image processing software. Neuronal branches were extracted by the neuron segmentation software SIGEN and stored in SWC format. **B.** Our revised segmentation scheme for a complex structured neuron. After the CLSM image was deconvolved by image processing software, masked image data were generated by tracing partial neuronal branches on the deconvolved image data. A partial branch image was obtained by applying AND image operation between the deconvolved image and the mask image. Another part of the neural branch image was generated by EOR image operation between the deconvolved image and the partial branch image. Neural segmentation software was applied on these two branch images independently. Whole neuron structure was reconstructed by combining these two reconstruction results.

To evaluate the performance of SIGEN, we compared segmentation results with those of other available segmentation tools. The BigNeuron project (*https://allenlnstitute.org/blgneuron/*) provides a huge collection of image data containing singly stained neurons along with corresponding gold standard segmentation results. The gold standard results were manually segmented from image data and were used as reference morphologies when comparing different algorithms. We used 40 samples from the package “p_changed7_janelia_flylight_part1”, which includes neurons of various shapes from fly brain imaged using confocal microscopes, as in our experiments. Parameter values for segmentation by SIGEN were as follows. Only those fragmented segments with volumes larger than 30 voxels were included in the segmentation (VT = 30). Further, fragmented segments whose distance from the main branch, after rounding to the nearest voxel, was less than 30 voxels (DT = 30), were connected to the reconstructed neuron. Segmentation results were evaluated using the DIADEM metric value (Gillette et al., 2011). DIADEM metric was developed and provided by the DIADEM Challenge project for evaluation of neural segmentation results by comparison to gold standard reconstructions. The DIADEM metric value ranges from 0 to 1 depending on the match condition of several features, such as node positions and branching state, with a value of 1 indicating a perfect match.

Metric values of segmentation results of an algorithm can vary largely depending on input image data. In particular, when multiple neuronal branches spread their dendrites into the same region, it can be extremely difficult to accurately extract their structural connectivity with automatic segmentation. To apply our segmentation software to such complicated neuronal structures, we propose a revised scheme incorporating a process of manually separating overlapping neurites (Fig. 1B). In our revised segmentation scheme, we traced the neurites manually to create a mask image for separating branches extending in the same region. For manual tracing, we used the 3D segmentation software, ITK-SNAP (Yushkevich et al., 2006; RRID:SCR_002010). The manual tracing operation was started from a clearly separated neuronal branch point. Neuronal segments belonging to the branch were determined by characteristics of the image, such as direction of intensity changes in a voxel cluster, characteristics of connectivity and bifurcation from the start point to peripherals. Of course, this process requires a certain level of skill and there is a possibility of human error, but at this time, it provides more reliable separation than by computer. Then, a partial neuronal image containing one of the overlapping branches was obtained by application of logical product (AND) image operation between the manual traced branching image and the deconvoluted neuron image. The image of the other overlapping branch was obtained by exclusive-or (EOR) image operation between the deconvoluted neuron image and the previously obtained partial neuron image. Since the EOR image operation results in the background value (0) when the values of voxels of the two images are the same, an image excluding the branches previously obtained from the original image is obtained. For the case where more neurites are densely concentrated in one region, the same procedure can be recursively applied by separating neurites into individual partial structures at each step.

The whole branching structure was obtained by conjunction of segmentation results of partial neurites. Finally, the segmented neuron structure was stored by the software in SWC and VTK formats. Thus, the results could be displayed and edited by other tools such as Vaa3D (Peng et al., 2010; RRID:SCR_002609), NeuTube (Feng et al., 2015), or ParaView (Ayachit, 2015; RRID:SCR_002516).

### 2.2 Honeybee neuron and its segmentation

Neuron images of DL-Int-1 in the honeybee brain were obtained experimentally. The details of intracellular recording and staining procedures were described in Ai et al. (2009; 2017) and we describe them here briefly. Honeybees (*Apis mellifera L.*) were reared in hives at the Fukuoka University campus. Forager worker bees, which collected pollen on their hind legs, were caught at the hive entrance and used in this study. In addition, newly emerged adults were isolated in a cage when they hatched from brood cells. Their ages were recorded in days, and only those less than 3 days old were used in our experiments. For the experiment, a bee was immobilized by cooling and was mounted in an acrylic chamber. The mounted bee was fed with 1 M sucrose solution and kept overnight in the dark with high humidity at 20°C. The head of the bee was fixed with wax. The frontal surface of the brain was exposed by cutting away a small rectangular window between the compound eyes. The glands and tracheal sheaths on top of the brain were removed. The mouthparts, including the mandibles, were cut off to expose and remove the esophagus. Small droplets of a honeybee physiological saline were applied to wash away the residue of the esophagus and to enhance electrical contact with a platinum ground electrode placed in the head capsule next to the brain.

Glass electrodes were filled at the tip with 3% Lucifer Yellow CH Dilithium salt (L0529, Sigma-Aldrich, Tokyo, Japan) dissolved in 100 mM KCl, yielding DC resistances in the range of 150 to 300 MO. We also used Dextran, Tetramethylrhodamine, 3000 MW, Anionic, Lysine Fixable (D3308, Thermo Fisher) and Alexa 647 hydrazide (A20502, Thermo Fisher, Tokyo, Japan) as dyes for the injection. The electrode was inserted into a region of the DL after the neural sheath and a small area of the brain’s neurilemma had been scratched. Electrical signals were amplified with an amplifier (MEZ 8301, Nihon Kohden, Tokyo, Japan) and displayed on an oscilloscope. After identifying DL-Int-1 by their unique response patterns to the vibration stimuli applied to the antenna, the fluorescent dyes were injected iontophoretically.

Thereafter, the brains were removed, fixed in 4% paraformaldehyde for 4 h at room temperature, and then rinsed in phosphate buffer solution, dehydrated, and cleared in methyl salicylate for subsequent observation. The cleared specimens containing intracellularly stained neurons were viewed from the posterior side of the brain under a CLSM (LSM 510, Carl Zeiss, Jena, Germany) with a Zeiss Plan-Apochromat 25×/0.8 NA oil lens objective (working distance 0.57 mm). Alexa 647 was excited by the 633-nm line of HeNe lasers, and Lucifer yellow was excited by the 488-nm line of an argon laser. More than 250 optical sections were taken at 1 μm thickness throughout the entire brain depth of each specimen. The image resolution of each section was 0.36 μm, and size was 1024 × 1024 pixels.

## 3 Results

### 3.1 Evaluation of semiautomatic segmentation scheme

To confirm the performance of our basic segmentation tool SIGEN, we applied it on samples provided by the BigNeuron project (https://glthub.com/BlgNeuron/Data/releases). Neuron images with gold standard segmentation data can be downloaded from the site. We used samples in the subset “p_packed7_janelia_flylight-part1”, of the gold166 package. A neuron with complex structure was reconstructed well by SIGEN, resulting in a DIADEM metric value of 0.850 (Fig. 2A). In contrast, the metric value was relatively low for a neuron with simpler structure (Fig. 2C). In this case, although thick and long branches were reconstructed well, SIGEN had difficulty tracing small dense neurites located in the lower right part of the image. However, experts also had difficulty manually tracing these dense arborizations, as seen from the gold standard results. The average value of the DIADEM metric value of SIGEN for 40 samples was 0.717 ± 0.163. Higher average values compared with other software and small values of variance indicate that neuronal morphology can be reconstructed stably by SIGEN (Fig. 3). This package contains images of various shapes of insect neurons. Because SIGEN was able to successfully handle such neuron images, the software is considered to be suitable for the reconstruction of neuronal morphology.

**Figure 2.**
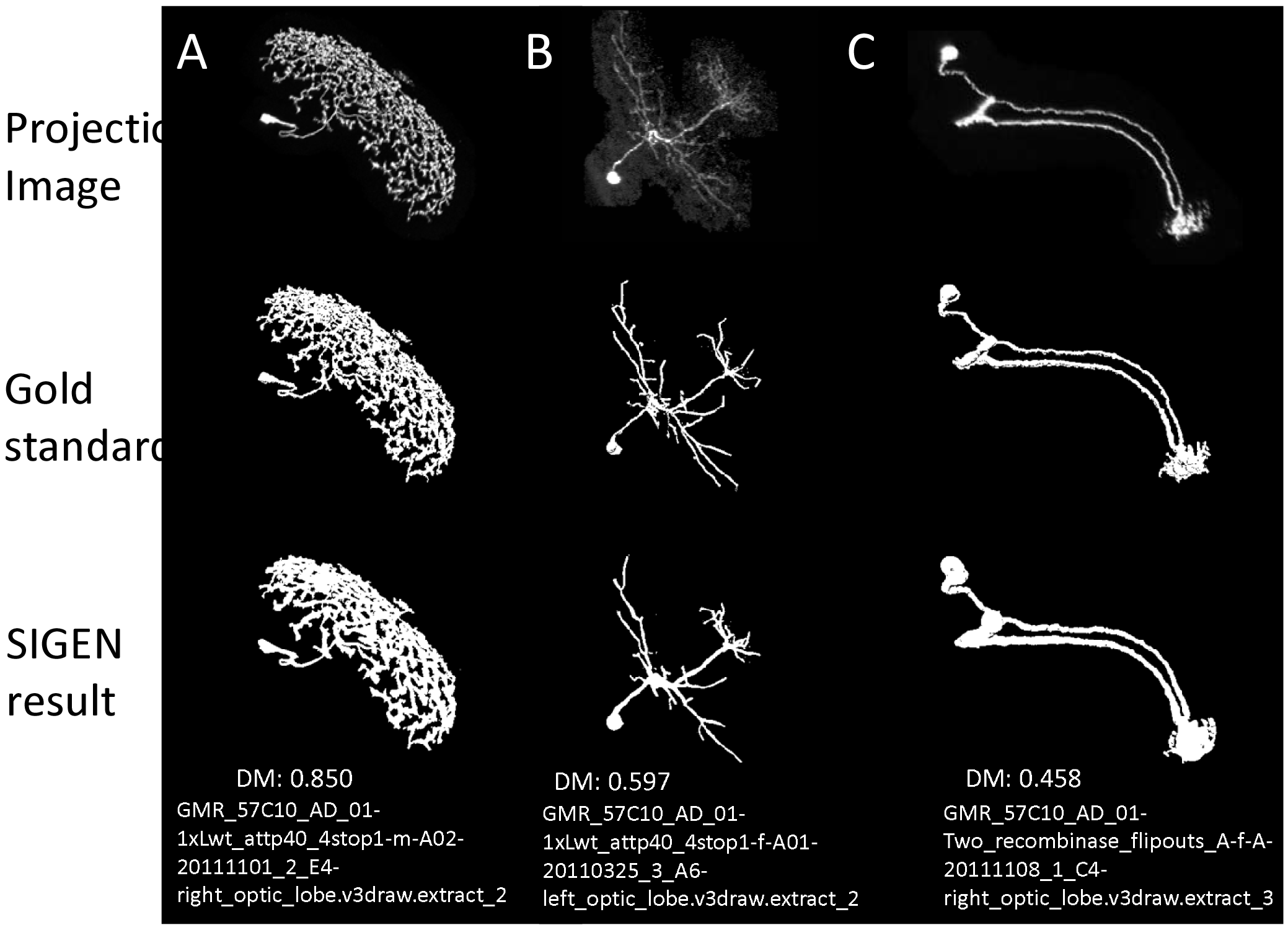
Three examples of segmentation by our original software, SIGEN, on samples provided by the BigNeuron project. *Top row*, 2D projected image data of neurons. *Middle* row, the manually segmented gold standard results, provided by BigNeuron. *Bottom* row, segmentation results by SIGEN. The difference between gold standard and automatic segmentation results was measured by the DIADEM metric, whose value ranges from 0 (completely unmatched) to 1 (perfectly matched). A neuron with a complex branching structure was reconstructed well, as shown in sample A (DIADEM value: 0.850). In contrast, the neuron shown in sample C was reconstructed with a low DIADEM metric value of 0.458, due to the presence of many tiny neurites that were difficult to extract, as was also seen in the gold standard results.

**Figure 3.**
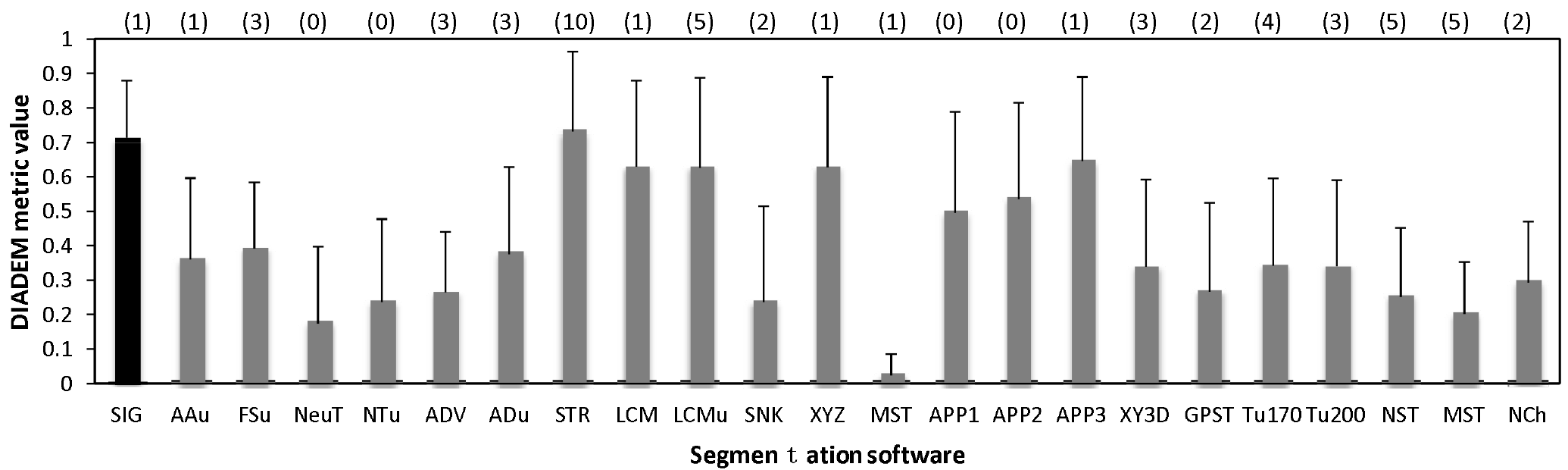
The average values of the DIADEM metric for 40 samples in the package “p_changed7_janelia_flylight_part1” of BigNeuron. Error bars indicate standard deviation, and the numbers in parentheses at the top of the graph are the number of samples for which no DIADEM value was obtained for the segmentation result. Abbreviations are segmentation software names: SIG, SIGEN; AAu, axis analyzer updated; FSu, fastmarching spanningtree updated; NeuT, neuTube; NTu, neuTube updated; ADV, Advantra; ADu, Advantra updated; STR, smartTracing; LCM, LCMboost; LCMu, LCMboost updated; SNK, snake; XYZ, All-path-pruning; MST, MOST; APP1, All-path-pruning1; APP2, All-path-pruning2; APP3, All-path-pruning3; XY3D, XY_3D_TreMap; GPST, NeuroGPSTree; Tu170, Tubularity model_S MST tracing th170; Tu200, Tubularity model S MST tracing th 200; NST, NeuroStalker; MST, MST Tracing; NCh, NeuronChaser updated.

In the image set used above, some of the samples have multiple neurites mixed in one region. In such cases, all software approaches failed to extract good reconstructions. For example, the neuron “GMR_57C10_AD_01-Two_recombinase_flipouts_A-f-A-20111108_1_C4-right_optic_lobe.v3draw.extract_3” has a primary neurite extending from the cell body with neurites spreading in a certain region (arrow in Fig. 4A). Furthermore, a secondary neurite from the dendrite runs along the primary neurite (Fig. 4A, B). It is important to grasp the state of extension of such a neuron accurately, and in the gold standard by manual extraction, the primary neurite folded near the secondary neurite was accurately extracted (Fig. 4C). However, in the reconstruction generated from our automatic segmentation software SIGEN (Fig. 1A), the midway and terminal points of the neurites were erroneously connected (compare Fig. 4C and Fig. 4D yellow arrows). The structure was significantly different from the gold standard, resulting in a DIADEM metric value of 0.488. The wrong interpretation of the neurite form was not limited to SIGEN. The DIADEM metric values for other automatic segmentation software were comparatively low, with a maximum of 0.544 and an average of 0.151. By applying our proposed method (Fig. 1B), neurites elongating in different directions were separated and extracted, and then synthesized (shown in red and green in Fig. 4B). It became possible to accurately extract the major cell structure, as shown by the large improvement of DIADEM metric value from 0.488 to 0.825 (Fig. 4E). Therefore, for cases where multiple dendrites extend to the same area, our proposed method is more effective than all compared automatic segmentation methods.

**Figure 4.**
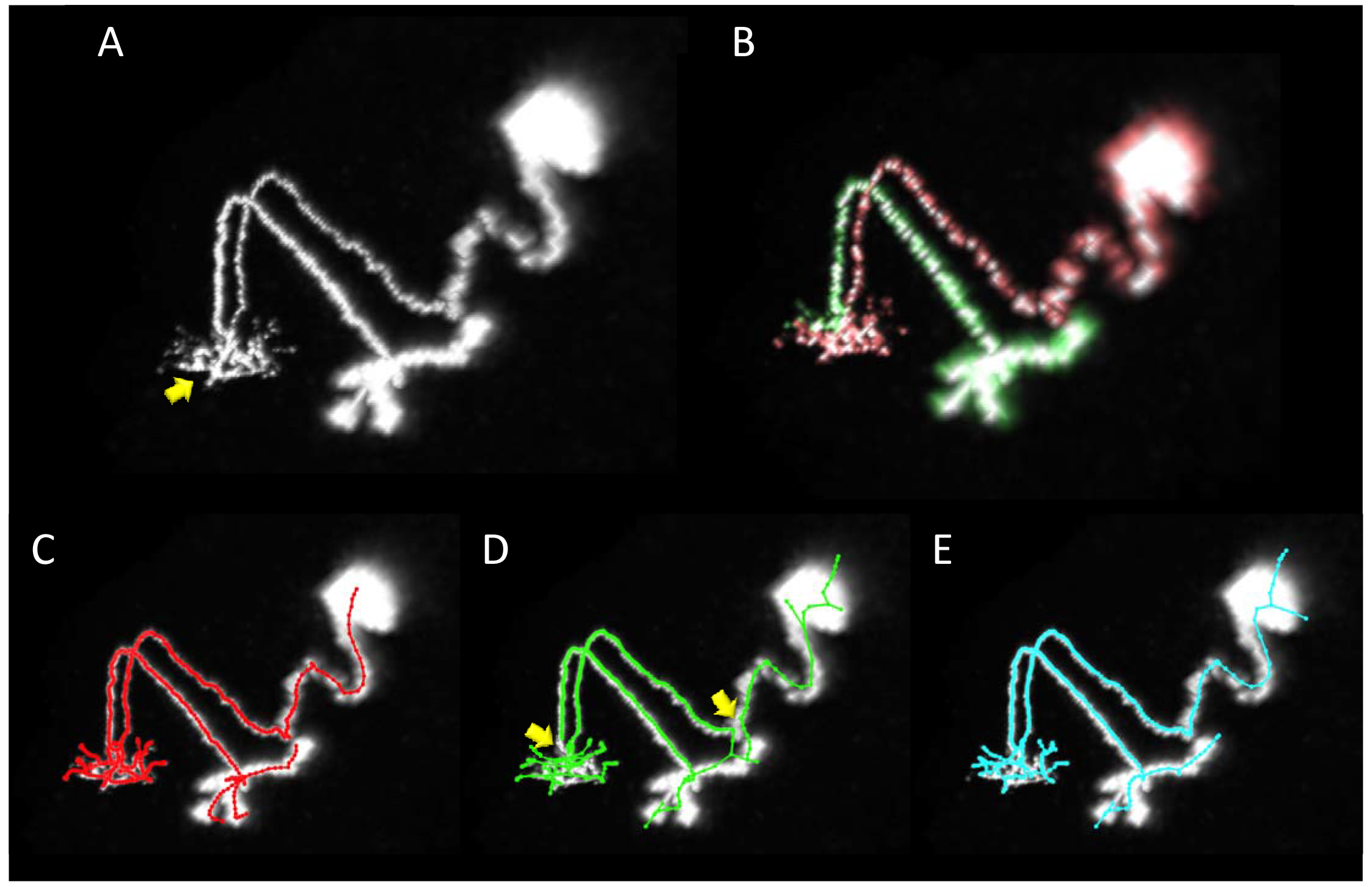
An example of segmentation of a neuron with complex structure. **A.** Projection image of the neuron. The primary neurite from the soma extends and arborizes in the region indicated by the yellow arrow. A secondary neurite runs parallel with the primary neurite. **B.** Two neurites (the soma and the primary neurite are shown in red, and the secondary neurite in green) were separated by manual operation in our revised scheme. **C.** The gold standard of segmentation. **D.** Segmentation result by our basic scheme (Fig. 1A). E. Segmentation result by our revised scheme (Fig. 1B). In the result of our basic scheme in D, there are segmentation errors on major connecting points of neurites (yellow arrows in C, D), most of which were rectified in the revised scheme in E (yellow arrows in E).

### 3.2 Segmentation of a vibration sensitive interneuron in honeybee brain

The soma of DL-Int-1 is located at the dorsal central body in the posterior protocerebral lobe (Fig. 5A) and extends a long branch into the DL with two main arborizations separated into dorsal and ventral regions (Fig. 5C). This neuron showed a tonic inhibitory response to a stimulus applied to the vibration sensitive region of the antenna (Fig. 5B). In this study, we focused on the morphology of DL-Int-1 dorsal and ventral arborizations in the DL (n_2_ and n_3_ in Fig. 5C, respectively). To obtain the whole arborizing image, separately captured confocal image stacks were stitched in 3D space using Fiji (Schindelin et al., 2012, Preibisch et al., 2008).

A neurite terminal was automatically selected as the start segment (root point) of segmentation by SIGEN. After the neuronal structure was obtained by our segmentation tool, we changed the root point on the branching node of the neurite (n_1_ in Fig. 5C) toward the soma. Shapes of neuronal branches were extracted well, but several connections among the branch segments were incorrect even at the major branching points, which connect to thick branches (white arrow in Fig. 6A). To correctly reconstruct the major neuronal structure, we applied our revised segmentation scheme. We generated a mask image for extracting the ventral branching part (Fig. 6B) from the whole neuron image. The manually-traced mask image of the ventral branch was applied by AND image operation on the deconvoluted image to obtain the ventral branch image (Fig. 6C). The dorsal part of the neuronal branch was extracted by EOR image operation between the deconvoluted image and the ventral branch image (Fig. 6D). Segmentation and reconstruction of neuron morphologies were generated from these ventral and dorsal images independently. The whole neuronal structure was reconstructed by connection of these two reconstructed results (Fig. 6E). Although the overall shape of neuronal branches obtained by the automatic (Fig. 1A) and revised schemes (Fig. 1B) was similar, connections of branches and distance from the root (start) point were quite different between the methods (Fig. 7). As shown in the enlarged view, the thick neurite extending to the ventral side branches into two neurites. One branch extends to the dorsal side and the other extends to the ventral side and branches again. Such a structure could be extracted correctly, as shown in Fig. 7B, because the revised scheme had grasped the whole structure in advance. In contrast, it could not be extracted by SIGEN alone, and it was disconnected just after branching like the result of Fig. 7A. By our revised segmentation scheme, we obtained reconstruction results from eight foragers (Fig. 8A) and seven age-controlled adults (Fig. 8B). Detailed analyses based on morphometric features, such as branching patterns, branch segment length and diameter, could be addressed in future research to assess morphological differences between ages.

**Figure 6.**
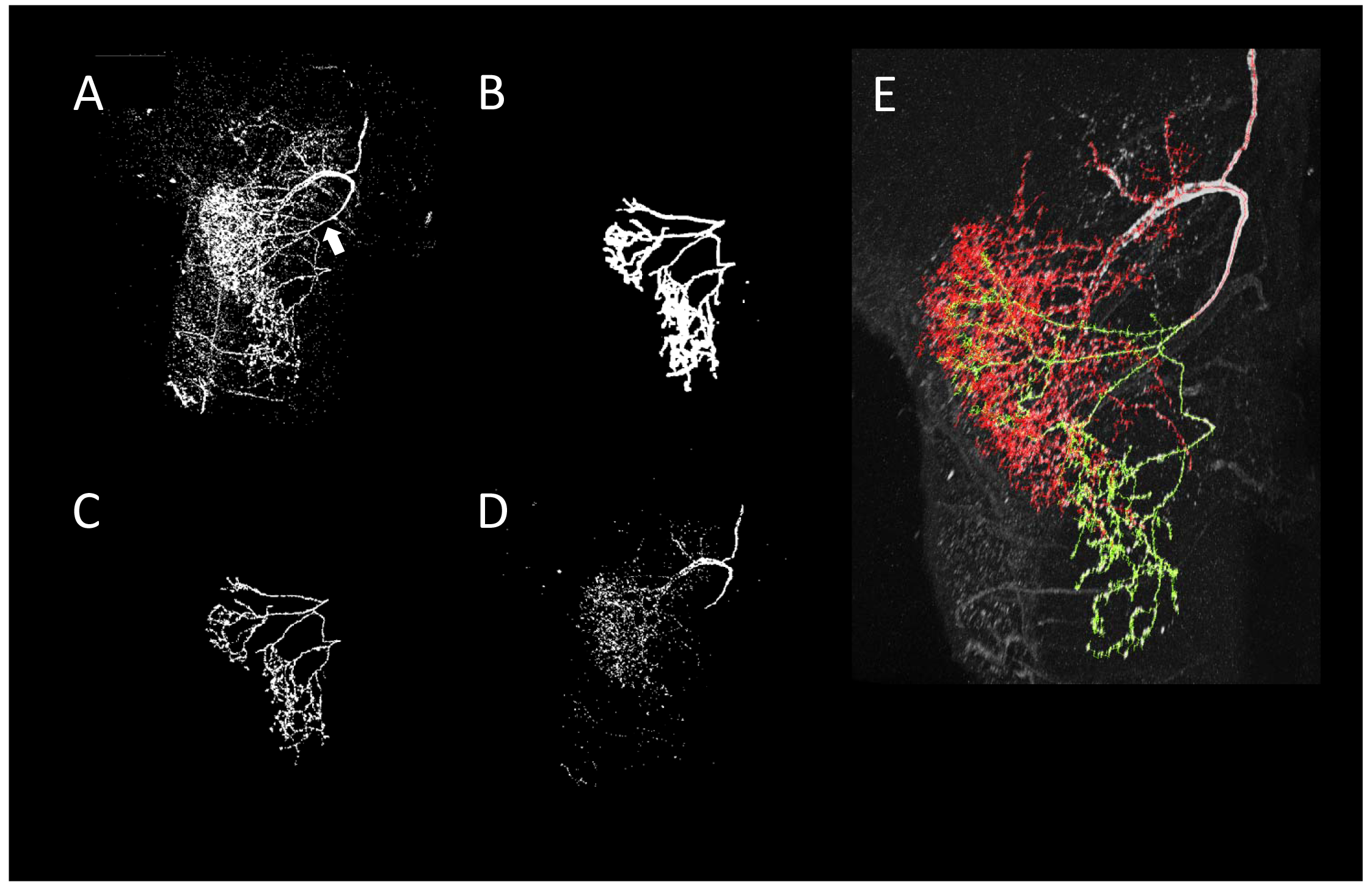
Reconstruction of neurite arborizations of DL-Int-1 by using our revised segmentation scheme. **A.** 2D projection image of DL-Int-1 in the dorsal lobe. The white arrow indicates the point that SIGEN could not extract the correct connection by our basic scheme. **B.** 2D projection image of a manually traced 3D mask filter for separation of ventral and dorsal branches. **C.** The extraction result of the ventral branches obtained by application of the mask filter (B) on the whole arborization image (A). **D.** The dorsal branches obtained by the EOR image calculation between whole branching (A) and ventral branches (C). **E.** Skeleton traces of reconstruction overlapped with the original 2D-projected image. Green and red skeleton traces show ventral and dorsal branches, respectively.

**Figure 7.**
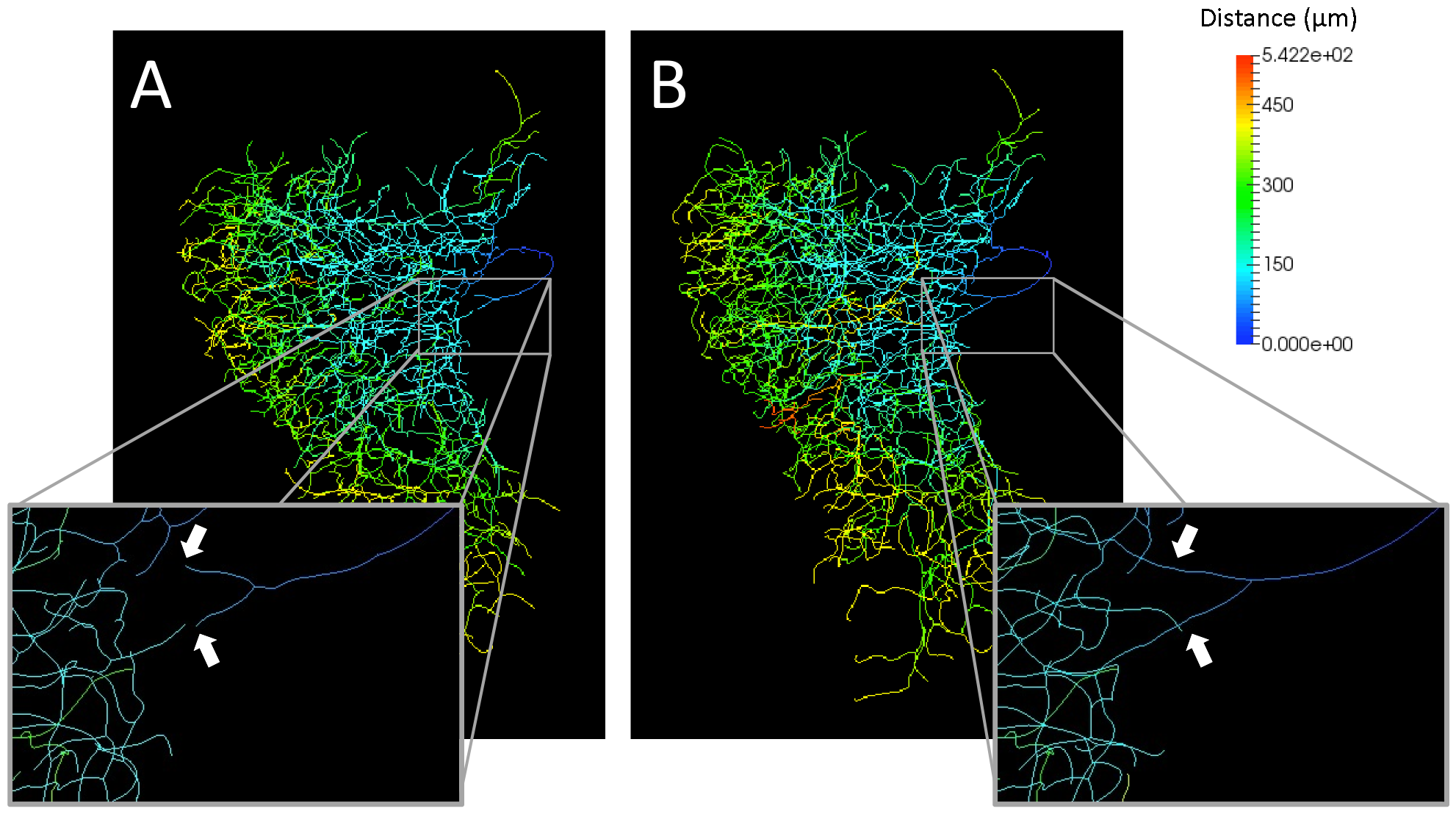
Comparison of distances from the root point on segmentation results by our basic (A) and revised (B) schemes. The distance from root points were indicated by pseudocolors. The enlarged views show many places where the connection is different between the basic scheme and revised scheme. In particular, the place where the incorrect connection state was obtained by the basic scheme is indicated by white arrows.

**Figure 8.**
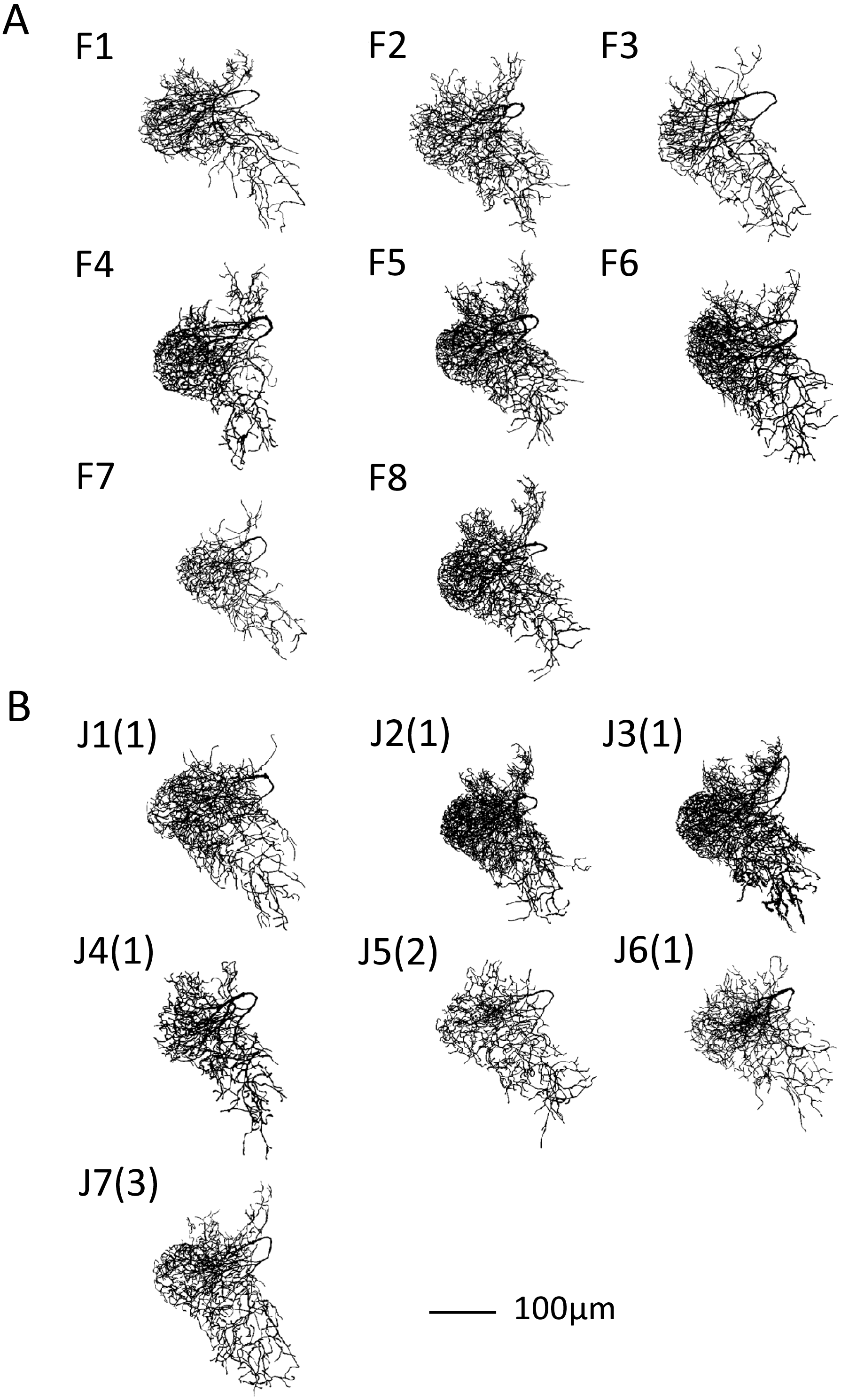
Segmentation results of DL-Int-1 for foragers, labeled by F+ID number, (A) and age-controlled adults, labeled by ID number with age (days) in parentheses (B).

## 4 Discussion

We applied the segmentation software SIGEN to CLSM images for constructing morphological models of neurons. It achieved high scores on benchmark tests of the BigNeuron samples. Its segmentation results demonstrated its reliability in extracting the major structure of a neuron without human bias. Various approaches to develop automatic neuron segmentation software are being undertaken in fields including neuromorphometric research and neural computation.

In this study, we also developed a segmentation scheme for neurons with complex arborizing patterns that performed better than the compared existing approaches. Preprocessing CLSM images was useful to separate the neuron image from the background. The main difficulty in extracting structures like that of DL-Int-1 was caused by multiple arborizations extending and overlapping in the same region. Hence, it was difficult to identify and connect them from fragmented images. We applied state-of-the-art automatic segmentation tools, such as App2 and SmartTracing (Chen et al., 2015), on these neurons. The reconstruction result involved many erroneous branches caused by noise and the background brain image. Although our revised scheme requires a user to conduct manual tracing to make a mask filter, it produced more accurate segmentation results for structurally complex neurons. Manual tracing can introduce a problem with reproducibility, but at the present time it provides a more accurate result than fully automated techniques. Moreover, in our scheme the manual step is represented by the mask filter, which can be saved as objective documentation of the intervention. A future goal is to have a fully automatic segmentation tool, which we attempt to develop based on the advanced segmentation software.

The DL-Int-1 interneuron is considered to play a key role in the primary auditory center for vibration signal processing. In this study, we have obtained more than 15 reconstructions of DL-Int-1 from foragers and newly emerged adults. Future research could apply detailed morphometric analyses to the segmentation results and evaluate the similarity and differences of neuronal morphologies with age.

## 5 Conclusion

Automatic or semiautomatic segmentation techniques have become quite useful for morphometric analysis of neurons. In this study, we combined manual masking and filtering with an existing software for semi-automated dendritic segmentation of interneurons with complex structures. The efforts to develop segmentation techniques will benefit from new algorithms for image processing and machine learning in the future. However, this process is ongoing because segmentation results depend on various factors such as neuron form and image quality. Therefore, there is a continuing need to develop better automatic segmentation software tools. Until such a new tool is obtained, we think that it is best to respond flexibly by a combination of automatic segmentation software and manual operation, as proposed here. By using our developed procedure, we could proceed to analyze age-and labor-dependent morphometric change of critical interneuron related with deciphering dance communication in honeybee.

## 6 Conflict of Interest

The authors declare that the research was conducted in the absence of any commercial or financial relationships that could be construed as a potential conflict of interest.

## 7 Author Contributions

HA, TW and HI designed this work. KK and HA performed experiments to acquire neuron image data. HI and AK conducted segmentations. HI drafted the manuscript and all authors reviewed and approved the final version of manuscript.

## 9 Acknowledgments

We thank Dr. Philipp Rautenberg for his valuable advice on this study and Mr. Sho Iizuka for his efforts on maintain and revising of our segmentation software.

